# Genetic signatures for lineage/sublineage classification of HPV16, 18, 52 and 58 variants

**DOI:** 10.1101/2020.07.30.229112

**Authors:** Zhihua Ou, Zigui Chen, Yanping Zhao, Haorong Lu, Wei Liu, Wangsheng Li, Chunyu Geng, Guohai Hu, Xiaman Wang, Peidi Ren, Na Liu, Shida Zhu, Ling Lu, Junhua Li

## Abstract

Increasing evidences indicate that high-risk HPV variants are heterogeneous in carcinogenicity and ethnic dispersion. In this work, we identified genetic signatures for convenient determination of lineage/sublineage of HPV16, 18, 52 and 58 variants. Using publicly available genomes, we found that E2 of HPV16, L2 of HPV18, L1 and LCR of HPV52, and L2, LCR and E1 of HPV58 contain the proper genetic signature for lineage/sublineage classification. Sets of hierarchical signature nucleotide positions (SNPs) were further confirmed for high accuracy (>98%) by classifying HPV genomes obtained from Chinese females, which included 117 HPV16 variants, 48 HPV18 variants 117 HPV52 variants and 89 HPV58 variants. The circulation of HPV variants posing higher cancer risk in Eastern China, such as HPV16 A4 and HPV58 A3, calls for continuous surveillance in this region. The marker genes and signature nucleotide positions may facilitate cost-effective diagnostic detections of HPV variants in clinical settings.

## Background

Human papillomaviruses (HPVs) are a heterogeneous group of double-stranded DNA viruses mainly infecting epithelial surfaces of human beings. Currently, more than 200 types of human papillomaviruses have been identified, with the majority clustering into a limited set of phylogenetic genera (e.g., Alpha-, Beta-, Gamma-, Mu- and Nu-PV) [1]. The genital high-risk carcinogenic HPV types (e.g., HPV types 16 and 18) causing cervical cancer are part of a monophyletic clade within the genus Alpha-PV [2,3]. Cervical cancer is the 4^th^ most common cancer in women [4], with more than 0.5 million new cases and 0.2 million deaths occur worldwide annually [5].

Distinct HPV types are defined based on the L1 open reading frame (ORF) genetic sequence, with dissimilarity at least 10% to all other characterized viruses as a “novel” HPV type. Isolates of the same HPV type are referred to variant lineages and sublineages based on the complete genome nucleotide sequences differing approximately 1%-10% and 0.5%-1%, respectively [6]. HPV16, 18, 52 and 58 have been found to be the top four prevalent high-risk HPVs among Chinese females [7– 10]. So far, multiple variant lineages and sublineages have been designated to HPV16 (A1-A4, B1-B4, C1-C4, D1-D4) [6,11,12], HPV18 (A1-A5, B1-B3, C) [6], HPV52 (A1-A2, B1-B2, C1-C2, D) [6] and HPV58 (A1-A3, B1-B2, C, D1-D2) [6]. HPV variants have different phenotypic characteristics including carcinogenicity and ethnic dispersion. For example, HPV16 A3 and A4 were linked with higher cancer risks in Asian populations comparing to the European prototype A1, while A4 and D displayed higher carcinogenesis in North Americans than A1 [12,13]. Moreover, HPV16 A1 and A2 tended to cause higher cancer risks in white Americans, and D2 and D3 in Latino Americans [11]. Similarly, HPV18 B/C, HPV52 B and HPV58 A3 might be linked with higher cancer risks than the other variants of the same type [14– 18].

Since infections with different HPV variants herald cancer risks differently, identification of lineage/sublineages will provide instructions on the triage and screening frequency of HPV-infected individuals. Herein, we used the publicly available genomes of HPV16, 18, 52 and 58 to pinpoint the marker genes and signature nucleotide positions for the determination of HPV variants, which would facilitate cost-effective diagnostic detections of HPV lineages and/or sublineages. Genomes of HPV16, 18, 52 and 58 types obtained from Chinese females were utilized for variant classification accuracy using the signature genes and sites.

## Methods

### Data preparation

Genome sequences for HPV16 (n=3,718), 18 (n=129), 52 (n=91) and 58 (n=172) were downloaded from NCBI nucleotide dataset by keyword search (Keyword: txid333760 for HPV16, txid333761 for HPV18, txid10618 for HPV52, txid10598 for HPV58; Species: Viruses; Molecular types: genomic DNA/RNA; Sequence type: Nucleotide; Release Date: from 0000/01/01 to 2019/07/25; Sequence length: from 7,000 to 8,500; accessed on 25 July 2019). Reference genomes [6,19] were retrieved from PaVE [20]. Only unique genomes with over 95% of the HPV genome length (after excluding ambiguous sites) were selected. Moreover, all genes (E1, E2, E4, E5, E6, E7, L1 and L2) and LCR (long control region) sequences of the selected genomes had >70% coverage of the corresponding gene/region complete length. The sequences were aligned with MAFFT v7.427 and manually scrutinized and edited with BioEdit v7.0.5. After exclusion of highly similar sequences, a total of 2,695 genomes (HPV16, n=2,385; HPV18, n=99; HPV52, n=77; HPV58, n=134; see **Supplementary Table 1**) were retained for downstream analysis.

**Table 1:** Accuracies of lineage/sublineage assignments using E1, E2, L1, L2 and LCR with verification dataset. Values indicate the percentage consistency in lineage/sublineage assignment using the corresponding sequences comparing against the assignment results using full genomes.

| Region | Sublineage |  |  |  | Lineage |  |  |  |
| --- | --- | --- | --- | --- | --- | --- | --- | --- |
|  | HPV16 | HPV18 | HPV52 | HPV58 | HPV16 | HPV18 | HPV52 | HPV58 |
| **E1** | 78.63 | 100 | 99.15 | 98.88 | 100 | 100 | 100 | 98.88 |
| **E2** | 100 | 100 | 99.15 | 73.03 | 100 | 100 | 100 | 73.03 |
| **L1** | 48.72 | 100 | 100 | 97.75 | 100 | 100 | 100 | 97.75 |
| **L2** | 49.57 | 100 | 97.44 | 98.88 | 100 | 100 | 100 | 98.88 |
| **LCR** | 94.02 | 100 | 100 | 98.88 | 100 | 100 | 100 | 100 |

### Phylogeny reconstruction and data visualization

Nucleotide substitution model test was conducted with IQ-TREE ModelFinder [21] and the best model identified was subsequently used for Maximum Likelihood (ML) phylogeny construction for HPV16, 18, 52 and 58 using the aforementioned datasets, with 1,000 ultrafast bootstrap pseudo-replications [22]. Lineage and sublineage assignments were conducted according to the classification criteria proposed by Burk et al. [6]. Representative phylogenies of HPV16, 18, 52 and 58 were reconstructed with mean intra- and inter-group sequence distances calculated using R package *seqinr* and in-house R scripts. Phylogeny and the associated data were visualized with *ggtree* package in R [23].

### Pairwise distance matrix comparison

Pairwise p-distances for HPV sequences were calculated with R package *ape*, with gaps deleted in a pairwise manner. Correlations between the DNA distance matrices of partial genomic sequences and the full genomes were analyzed using Mantel test with the R package *vegan* [24].

### Determination of signature nucleotide positions

All sequences over 95% of the reference complete genomes were used to identify lineage- and sublineage-specific signature nucleotide positions as marker sites for variant classification. Positions with over 10% gaps or displaying a conservation rate of over 98% across the alignments were excluded. All signature nucleotide positions were highly conserved in 99% of the sequences in the corresponding lineage/sublineage or genetic cluster. The HPV reference genomes used were downloaded from NCBI except for HPV16: HPV16, K02718 (downloaded from PaVE [20]); HPV18, AY262282; HPV52, X74481; HPV58, D90400. The distribution of the signature positions along the HPV genomes were further summarized based on a sliding window size of 1000bp and a step size of 500bp.

### HPV-positive cervical samples from Chinese women

Exfoliated cervical cells were obtained from women participating in National Cervical Cancer Screening Program in Eastern China, including Anhui, Jiangsu, Shandong and Guangdong provinces. HPV DNA detections were conducted with BGI SeqHPV Kit (BGI-Shenzhen, China) [25,26]. The majority of subjects involving in this large-scale screening program displayed no clinical illness or slight inflammation in histopathological examination. Only samples from participants who consented to donate their residual sample for microbial investigation were selected, with their personal data anonymized. A total of 347 participants were recruited in this study. All the participants aged from 30 to 67 years old, with a median age of 48.

### HPV genome sequencing and assembly

Probes covering the complete genomes of 18 HPV types (HPV6, 11, 16, 18, 31, 33, 35, 39, 45, 51, 52, 56, 58, 59, 66, 68, 69 and 82) were designed by MyGenostics [27]. The extracted DNA was sheared to fragments around 250 bp in length, adapter-added, hybridized with probes at 65°C for 24h, and washed to remove uncaptured fragments. The DNA library was sequenced using a BGISEQ-500 platform for paired-end 100bp reads (BGI-Shenzhen, China). Following demultiplexing, raw reads were trimmed with fastp [28] and deduplicated with BBMap (https://sourceforge.net/projects/bbmap/). Quality-filtered reads were mapped to HPV reference genomes with BWA alignment tool [29]. Reads with both ends aligned to HPV16, 18, 52 and 58 were extracted and subject to *de novo* assembly with SPAdes 3.12.0 [30].

### Verification of signature genes and nucleotide positions

HPV16 (n=120), 18 (n=48), 52 (n=120) and 58 (n=93) genomes from Chinese females generated in this work were used to verify the hierarchical signature nucleotide positions for variant lineage/sublineage classification. The designations were conducted based on ML tree topologies and signature nucleotide position mapping algorithms. In brief, the genomes combining with the reference sequences (**Supplementary Table 2**) were aligned with MAFFT v7.427 and constructed for ML trees using IQ-TREE with 1,000 ultrafast bootstrap implementations [21,22]. The phylogenetic trees inferred from the ORFs and LCR were reconstructed following the same procedure, and the subsequent classification results were compared against those defined by complete genome data. R scripts were developed in-house to map the sequences against the signature nucleotide position sets to define variant lineages and sublineages.

### Ethical statement

This study was reviewed and approved by the Institutional Review Board of Beijing Genomics Institute, Shenzhen, China (BGI-R071-1-T1 & BGI-R071-1-T2). All the participants consented to the donation of their exfoliated cell samples and anonymized associated data for research purposes.

### Data availability

The data that support the findings of this study have been deposited into CNSA (CNGB Sequence Archive) of CNGBdb with accession number CNP0001117 (https://db.cngb.org/cnsa/).

## Results

### Signature genes for sublineage/lineage identification

In order to pinpoint marker genes for HPV16, 18, 52 and 58 variant classification, we compared the consistency between the complete genomes, individual ORF/region (E1, E2, E4, E5, E6, E7, L1, L2, LCR), and the concatenated regions (4R: E1+E2+L1+L2, 5R: E1+E2+L1+L2+LCR, 8R: E1+E2+E4+E5+E6+E7+L1+L2, and 9R: E1+E2+E4+E5+E6+E7+L1+L2+LCR) regarding to sequence diversity and lineage/sublineage assignment.

### Sequence diversity comparison

All the partial genomic regions showed a positive correlation with full genome sequences based on pairwise distance matrix comparison (p<0.05) but yielded different correlation coefficient values. The distance matrices of the concatenated partial genomes usually display a high correlation with that of the full genome, with correlation coefficients ranging from 0.957 to 0.996 (**Figure 1**). For individual genomic regions, E2 of HPV16, L2 of HPV18, and E1 of HPV58 displayed the highest correlation with its full genome. For HPV52, the distance matrices of L1 and LCR displayed similar correlation with that of full genome depending on the nucleotide substitution model utilized.

**Figure 1:**
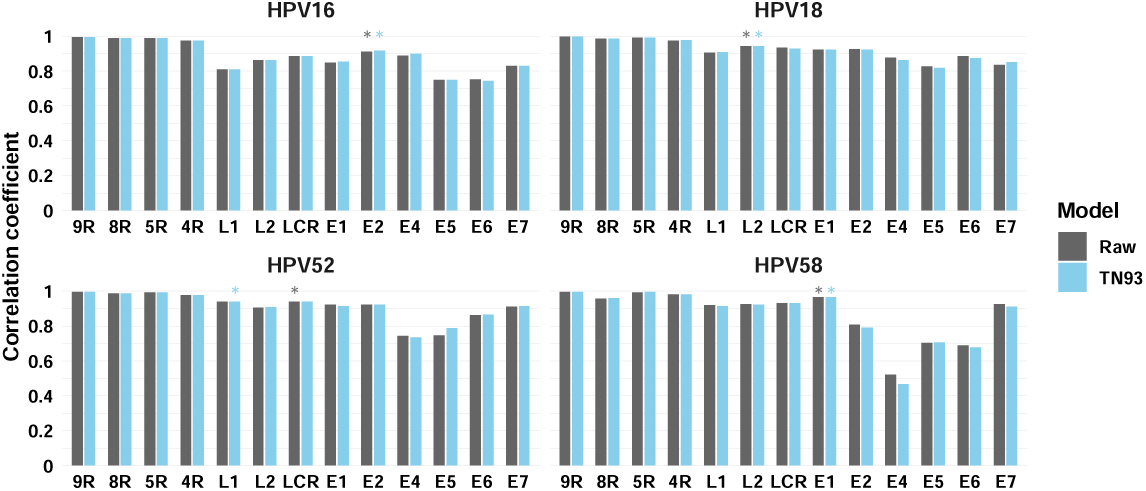
Correlation of the genetic distance matrix between full genome and partial genomic sequences of HPV16, 18, 52 and 58. Raw pairwise p-distances and genetic distances based on the TN93 model for HPV sequences were calculated using R package *ape*, with gaps deleted in a pairwise manner. Correlations between the DNA distance matrices of partial genomic sequences and the full genomes were explored using Mantel test with Spearman correlation method. Asterisks indicate the individual gene that showed the highest correlation with full genome under the corresponding substitution model for each HPV type. Abbreviations: FG, full genome; LCR, long control region; 4R, partial genome concatenated with 4 genetic regions: E1+E2+L2+L1; 5R, partial genome concatenated with 5 genetic regions: E1+E2+L2+L1+LCR; 8R, partial genome concatenated with 8 genetic regions: E6+E7+E4+E5+E1+E2+L1+L2; 9R, partial genome concatenated with 9 genetic regions: E6+E7+E4+E5+E1+E2+L1+L2+LCR.

### Lineage/sublineage classification

Phylogeny of each genomic region for lineage/sublineage classification was reconstructed with the same setting for each HPV type. The assignments based on partial genomes (e.g., 4R, 5R, 8R and 9R) were 100% consistent with those by complete genomes for HPV18, 52 and 58 (n<150). For HPV16, the classification consistencies of partial genomes were around 99.8%, which may be due to the relatively abundant dataset (n=2385) and the variation in the non-coding regions of this type (**Figure 2**). We also found high accuracy of variant classification inferred from distinct ORF/region including HPV16 E2 (98.49%), HPV18 L2 (100%), HPV52 L1 (100%) and LCR (100%), and HPV58 L2 (100%), LCR (100%) and E1 (100%). The phylogenetic topologies inferred from E4, E5, E6 and E7 genes were severely distorted, given the limited variation these genes contain. Although some short ORFs, e.g., HPV18 E4 and E5, HPV52 E6 and E7, achieved lineage/sublineage classification results with high accuracies in our study, the assignment process relied heavily on the abundance of references and experience, and may be prone to human subjectivity and visual error. Therefore, the short ORFs (E4, E5, E6, E7) may be not suitable for phylogenetic classification.

**Figure 2:**
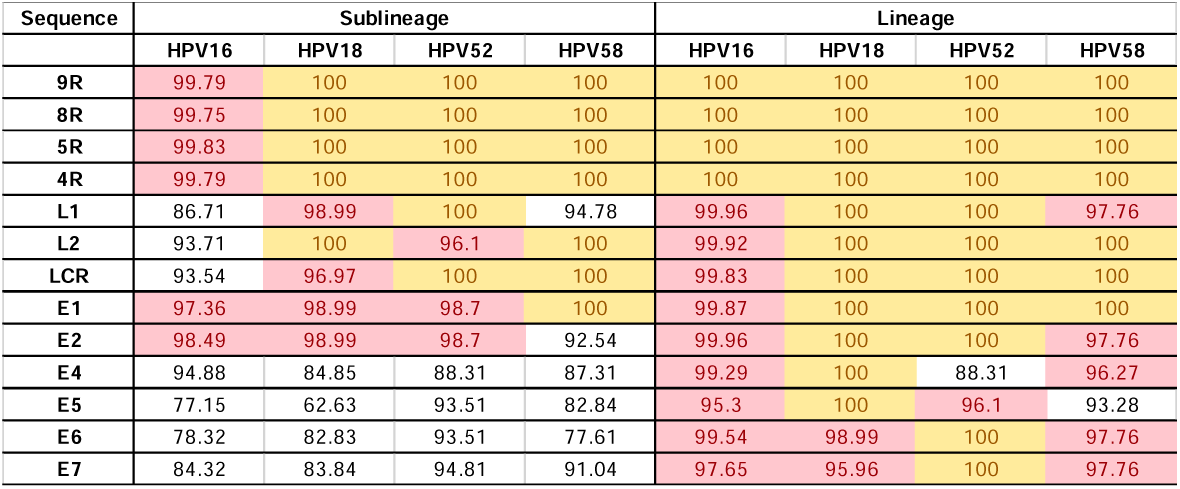
Accuracies of lineage/sublineage assignments using different part of the genomes. Values indicate the percentage consistency in lineage/sublineage assignment using the corresponding genomic regions comparing against the assignment results using full genomes. Cell colors: yellow, 100%; red: 95∼100%; white, <95%. Abbreviations are the same as **Figure 1**.

### Signature nucleotide positions for lineage/sublineage assignment

Since HPVs have a relatively low mutation rate [31] and previous reports have shown certain positions to be population- or lineage-specific, we sought to identify signature nucleotide positions with lineage and/or sublineage fixation across the complete genome. Because no exclusive single nucleotide positions were determinable for HPV16, 18, 52 and 58 variants, we used tree topologies and a hierarchical manner to identify lineage- and/or sublineage-specific nucleotide sites and patterns.

Certain sublineages, including HPV16 A1-3, B2-4, C1-4 and D1-4, HPV18 A1-4 and B1-3, and HPV52 A1-2 were merged together because of the limited sequence variation or inadequate distances these clusters contained (**Supplementary Figure 1**). A total of 79, 133, 161 and 123 signature nucleotide positions were characterized for HPV16, 18, 52 and 58, respectively (**Figure 4** and **Supplementary Table 3**). At lineage level of HPV16, 35 sites were able to discriminate lineage A from lineage B/C/D, followed by 13 sites further discriminating B from C/D, and 17 discriminating C from D. At the sublineage level of HPV16, we found 12 positions for the discrimination of A1-3 and A4, and 2 for B1 and B2-4. The hierarchical structure of signature nucleotide positions for HPV18, 52, and 58 can be interpreted similarly (**Figure 3**).

**Figure 3.**
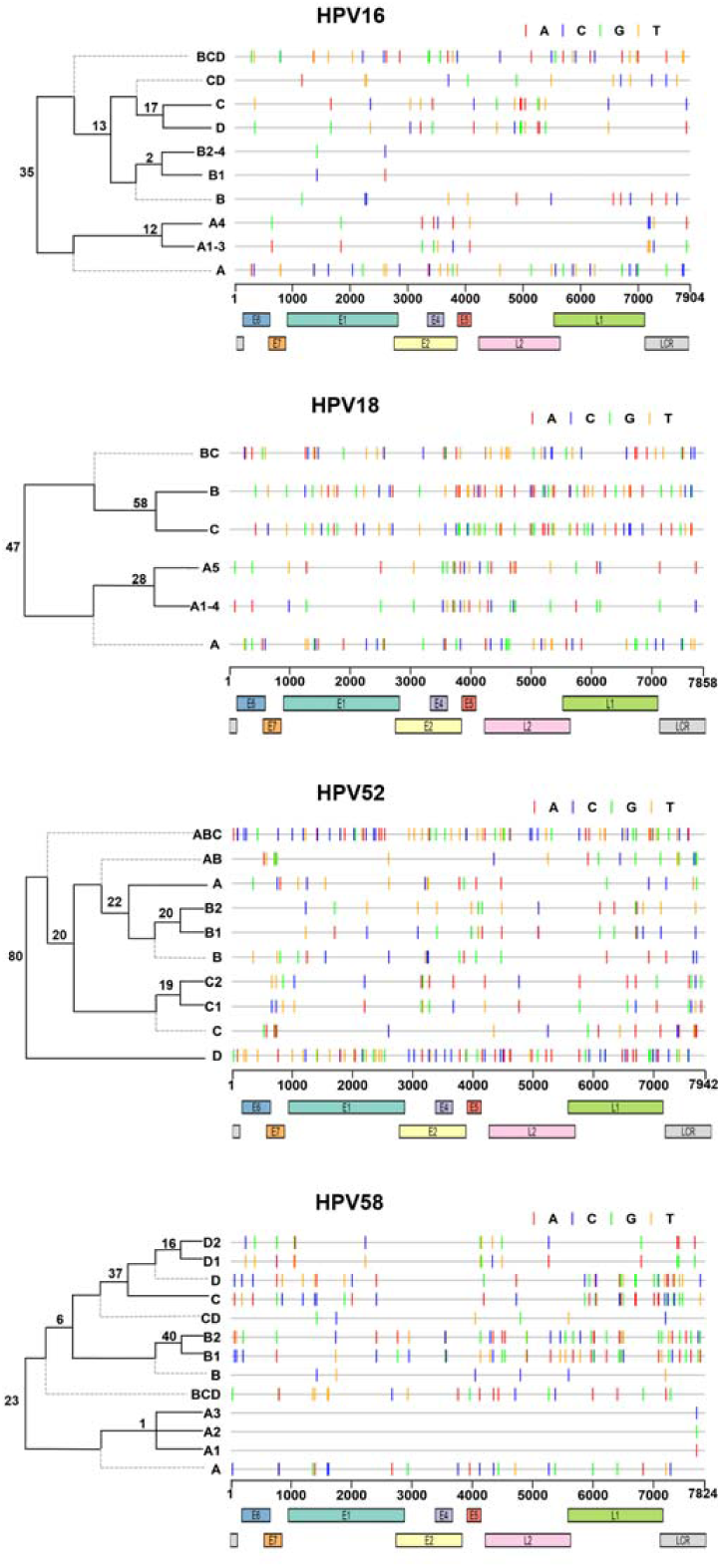
Hierarchical discrimination of HPV lineages and sublineages with signature nucleotide positions. The number of nucleotide positions determined for discriminating variant lineages and/or sublineages are displayed on the common node of two lineage/sublineage level. For example, in the panel of HPV16, 35 is the number of signature sites to distinguish lineage A and BCD, and 12 to distinguish A1-3 and A4. The hierarchical structure was not scaled to genetic distances. The signature positions were color-labelled for each classification level based on the genomic organization of the reference genome of each HPV type, with the ORF/LCR locations indicated by color bars.

**Figure 4.**
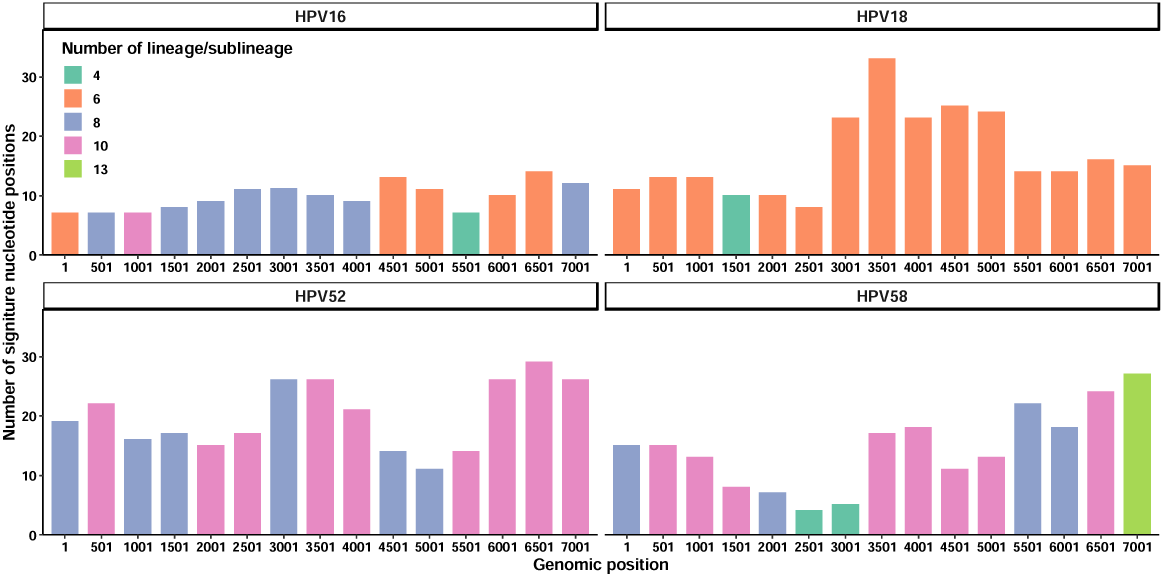
Genetic hotspots for signature nucleotide positions of HPV16, 18, 52 and 58 types. Using a sliding window size of 1000bp and a step size of 500bp, distribution of the signature nucleotide positions identified for the four HPV types were summarized. The X axis indicates the start position for each sliding window. The Y axis indicates the number of distinct signature positions for each window based on the corresponding reference genomes. Color code indicates the number of lineage/sublineage identifiable within each sliding window. The color legend applies to all types.

Using a sliding window size of 1000bp and a step size of 500bp, we explored the distribution of the signature sites along the viral genomes. Results showed that the 1001^st^-2000^th^ genomic positions of HPV16 (the 5’ terminal region of E1), the 3501^st^- 4500^th^ of HPV18 (flanking E2, E4, E5 and L2), the 6501^st^-7500^th^ of HPV52 (the 3’ terminal region of L1) and the 7001^st^-7824^th^ of HPV58 (LCR) may contain the most sufficient signature sites to distinguish all the hierarchical levels for each HPV type. (**Figure 4**).

### Genetic diversity of verification dataset

To verify the signature of marker genes and sites in variant classification, we accessed HPV16, 18, 52 and 58 genomes from Chinese women who participated in the National Cervical Cancer Screening Program. Most of the participants were from Eastern China. All sequences are >95% coverage in size of the complete genomes. Based on tree topologies and distance threshold, 116 out of 117 HPV16 sequences were unambiguously assigned as lineages A (A1-2=25, A3=24, A4=66) and D (D3=1) (**Figure 5, Supplementary Table 4**). The high prevalence of HPV16 A4 variants in Eastern Chinese women was consistent with previous reports in Eastern Asian [18]. HPV16 A4 was also linked with higher cancer risks in Asian populations than other variants [12,13]. All HPV18 genomes belonged to lineage A (A1=37, A3=1, A4=10) (**Figure 5, Supplementary Table 4**), which were consistent with previous reports on HPV18 sublineage distribution in China [32,33]. The majority of HPV52 and HPV58 sequences were lineages B (111/117, 94.9%) and A (88/89, 98.9%), respectively, that were further divided into HPV52 B2 (n=111) and HPV58 A1 (n=51), A2 (n=25) and A3 (n=12) (**Figure 5, Supplementary Table 4**). Rest of HPV52 sequences belonged to A1 (n=2), C2 (n=3) and D (n=1), and HPV58 to B2 (n=1). HPV52 lineages B and C were common in Asian countries, with B the most prevalent in China [34–36]. However, the cancer risk of HPV52 B variants was reported to be lower than lineage C [16]. Lineage A of HPV58 was found to be globally distributed and was the most prevalent in Asian females [16,33,35,37,38]. Moreover, HPV58 sublineage A3 might pose higher cancer risk than other variants [17,18]. The prevalence of HPV variants associated with higher cancer risk (e.g., HPV16 A4, HPV58 A3) in Eastern China called for continuous surveillance on female populations in this region.

**Figure 5.**
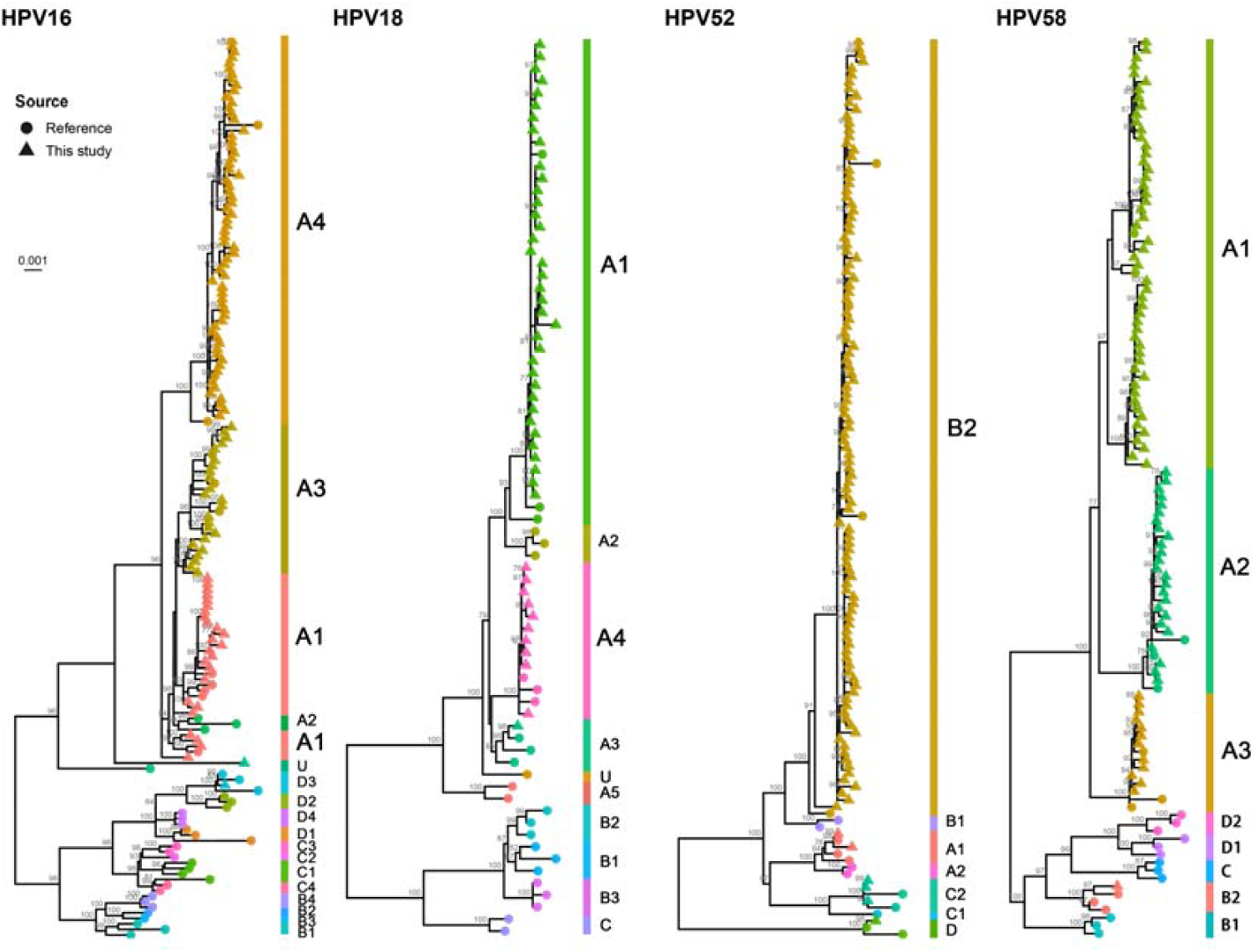
ML phylogenies of HPV16, 18, 52 and 58 sequenced in this study. The ML phylogenies were constructed with full genomes using IQ-TREE, implementing 1000 ultra-fast bootstrap tests. Bootstrap values of over 70 were displayed on nodes. Tips labelled by solid circles are reference genomes obtained from NCBI, while those labelled by triangles were sequenced in this study. Lineage/sublineages are indicated by colors and the name for each lineage/sublineage is presented with the same color. U: Undefined.

### Verification of HPV lineage classifications with signature regions and nucleotide positions

Using the assembled HPV sequences from Eastern Chinese women, we further confirmed that HPV16 E2, HPV18 L2, HPV52 L1 and LCR, and HPV58 LCR contained sufficient variation information to assign the target genes/region with proper lineages and/or sublineages, consistent with the classification by the complete genomes (**Table 1, Supplementary Table 4**). In addition, HPV58 E1 and L2 reached 98.88% accuracy in variant classification, except for one sequence with ambiguous assignment. Classification results by signature nucleotide positions also showed high consistency with those by the complete genomes, with 100% and >98.9% accuracies in variant classification at lineage and sublineage levels, respectively. It’s worth noting that one HPV58 A2 sequence based on the complete genome assignment was grouped to A1 since only one variation was discriminative for A1, A2 and A3. Hence, mutations at certain individual position may affect the accuracy of sublineage assignment.

## Discussion

It has been recommended to use the complete genomes to identify HPV variant lineages and sublineages [6]. However, ambiguous assignment may arise when the complete genomic sequences are not available in clinical settings or in developing areas. Lineage fixation of genetic changes in one gene/region highly correlated with other changes within genomes from the same lineage and sublineage is observed throughout HPVs and may represent adapted variations in natural selection with different phenotypic characteristics and carcinogenicity. In this study, we sought to identify genes and sites with lineage- and sublineage-specific significance for HPV variant characterization in large-scale screening program. Our data indicated that marker genes (e.g., HPV16 E2, HPV18 L2) as well as signature nucleotide positions proved high accuracy in lineage/sublineage classification, which was further verified by assembled HPV sequences from Eastern Chinese women.

Hotspots of the genetic signature position may be chosen as the target regions for the developing of rapid classification assays, such as the 1001^st^-2000^th^ genomic positions of HPV16, the 3501^st^-4500^th^ of HPV18, the 6501^st^-7500^th^ of HPV52 and the 7001^st^-7824^th^ of HPV58 (**Figure 3, Figure 4, Supplementary Table 3**). Because most detection methods, such as qPCR, utilize short genomic regions as detection targets, we recommend using multiple genetic regions to achieve optimal detection accuracy. Due to the uneven distribution of the genetic signature positions, single regions may not be able to provide high-resolution classification. For example, while the 1001^st^-2000^th^ positions of HPV16 may be able to distinguish all the classification levels, this region contains much less information for the separation of Ax sublineages than the 3001^th^-4000^th^ region. Therefore, selection of multiple genetic regions pertinent to the local HPV diversity and detection methods such as multiplex-PCR may facilitate the cost-effective classification of HPVs to lineage/sublineage levels, which would help promote large-scale epidemiological study on the carcinogenesis of HPV variants.

This study has several limitations: 1) Though we have already gathered all available genomes from public database, the global diversity of HPVs may not be comprehensively covered. The scarcity of high-quality full genomes for certain lineages and sublineages, such as HPV16 lineage C and D, hinders the exploration of cluster-specific genetic signatures. 2) Due to the limited sample sizes of this study, the genetic diversities of high-risk HPVs in China remain to be explored. 3) Because the clinical information of the surveyed sequences was not available in this work, the association between disease statuses and specific variants/variations in China remains elusive. Such defects call for genetic studies on HPV-infected individuals from more diverse geographic regions with sufficient clinical records to enhance our understandings of specific variants that may pose significant effects on cervical health.

The sublineage distributions of the four HPV types in China generated in this study were consistent with previous reports [16,33–40]. However, with the intensification of globalization, the genetic diversity of HPV in China and other parts of the planet remains dynamic and requires continuous surveillance. The genetic signatures characterized by this study may provide valuable references for the design of cheap and fast detection assay to classify the four high-risk HPV types in Eastern China. Nevertheless, high-quality genomes were still scarce for many HPV types except HPV16. With the extensive application of whole genome sequencing in HPV research, the classification power using signature regions or nucleotide positions could be further increased.

## Supplementary Data

**Supplementary Table 1: Genomes of HPV16, 18, 52, 58 downloaded from public database**.

**Supplementary Table 2: Reference HPV complete genomes selected for the four HPV types**.

**Supplementary Table 3: Hierarchical SNPs for lineage/sublineage classification of HPV16, 18, 52 and 58**.

**Supplementary Table 4: Classification of the HPV16, 18, 52, 58 genomes generated by this study**.

**Supplementary Figure 1: Representative phylogenies of HPV16, 18, 52 and 58**. Using publicly available complete genomes, ML trees were reconstructed for the four HPV types and each sequence was classified to sublineage level based on tree topologies. The representative phylogenies were reconstructed based on mean intra- and inter-group percentage differences of clades, which are simultaneously displayed on the right panel of each tree. The number of HPV genomes in each clade is displayed in tip labels. The sizes of black circles on tree tips indicate the relative sequence abundance of the corresponding clade. Abbreviations: Ax and Cx, the strains belonged to lineage A and C, but could not be assigned to any existing sublineages; U, (i.e., Undefined), the strains were not assigned to any existing lineage. Number of genomes used: HPV16, n=2,385; HPV18, n=99; HPV52, n=77; HPV58, n=134.

## Notes

### Author contributions

J.L., Z.O. and Z.C. designed the study. J.L., N.L., L.L., S.Z. and X.W. coordinated sample collection. H.L., W.L., G.H., C.G. and P.R. conducted viral genome sequencing. Z.O. and W.L. conducted data analysis. Z.O. wrote the manuscript. Z.C., J.L., Y.Z. and L.L. provided critical revision of the manuscript.

### Information of corresponding author

Junhua Li; PhD; Tel: 0086-13929566296; Affiliations: Shenzhen Key Laboratory of Unknown Pathogen Identification, BGI-Shenzhen, Shenzhen 518083, China & School of Biology and Biological Engineering, South China University of Technology, Guangzhou, China; Address: China National GeneBank, Dapeng New District, Shenzhen 518120, China.

### Related Meetings

Part of the lineage/sublineage classification results has been presented in EUROGIN 2019 (December 4-7, 2019, Monaco). Poster Number: #0397; Title: Preliminary Analysis on the Genetic Diversities of High-risk Human Papillomaviruses in Chinese Women.

## Acknowledgments

We warmly thank Dr. Houshun Zhu, Mr. Xianchu Cai, Ms. Jieyao Yu, Ms. Wei Zhou, Mr. Hongcheng Zhou, Miss. Yi Wang and Miss. Di Wu for their assistance in viral genome sequencing and dataset preparation. We thank China National GeneBank for providing sequencing service for this project. Thanks also to all the authors for contributing HPV genomes to NCBI GenBank. Last but not the least, the authors are grateful to the inspirational communication from Miss Shanshan Mo, Miss Feiyun Ou, Mr. Geer Xi and Miss Xiaobai Zhong.

---

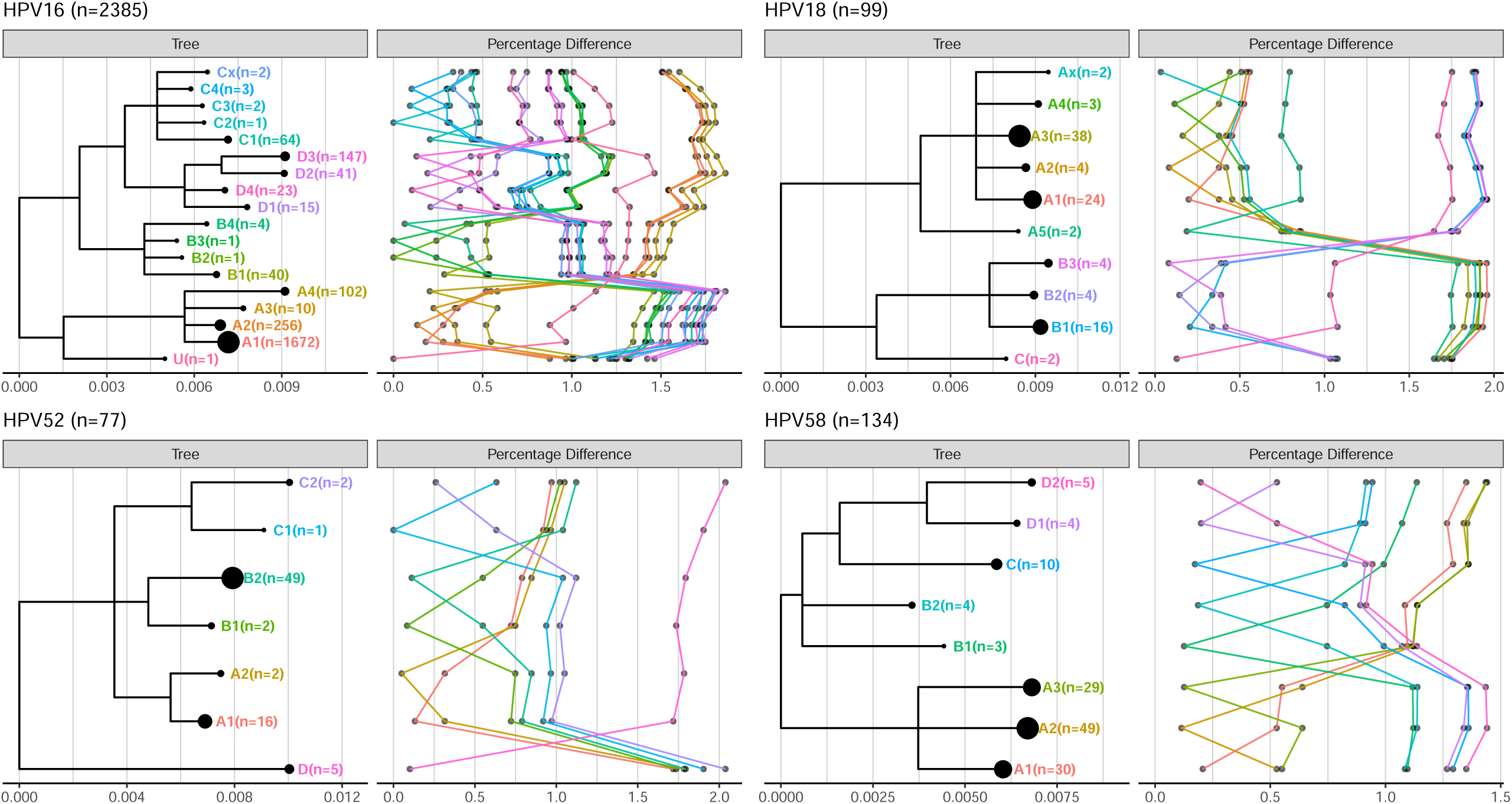

